# Drought-associated genes exhibit high constitutive expression in *Quercus douglasii*, a drought tolerant California oak

**DOI:** 10.1101/2025.07.07.663554

**Authors:** Stephanie E. Steele, Lily D. Peck, Victoria L. Sork

## Abstract

Drought stress is a strong selective pressure for all plant species. Plants can respond to water shortage through various strategies that confer drought tolerance, but with a potential cost for growth. These strategies may be plastic responses that occur with the onset of stress or may comprise continuously-expressed (constitutive) traits regardless of water availability. Here, we used RNA-Seq to characterize transcriptional responses to dehydration in seedlings of a drought tolerant oak, *Quercus douglasii*, from a population in the Sierra Nevada Foothills in California. In the greenhouse, we subjected 24 seedlings from six maternal families to dry-down or well-watered treatments and prepared RNA libraries from tissue collected before and after each treatment (48 libraries). Our goals were to characterize the pattern of up- and down-regulated genes in response to dehydration and to assess the extent to which this drought tolerant species shows differential versus constitutive expression as a drought response strategy. We identified few differentially expressed genes in response to dehydration. Up-regulated genes were related to known drought response functions, while down-regulated genes were enriched for gene ontology terms related to growth and carbohydrate metabolism. We discovered high constitutive expression of many putatively drought-responsive genes that had been found to exhibit gene expression plasticity in a drought sensitive oak, which a novel finding for drought tolerance strategies in tree species. We conclude that the drought tolerant strategy of *Q. douglasii* incorporates high constitutive expression levels of drought-responsive genes, as well as some plasticity in its response once environmental stress is experienced.

## Introduction

Drought is a major source of stress for plants and a strong selection pressure on traits that minimize that stress because intermittent availability of water interferes with core functions. Predicted increases in temperature and decreases in precipitation threaten forests globally (Allen et al. 2010), and large-scale regional die-offs due to drought have already been documented for many tree species (Potter 2016; Choat et al. 2018; Restaino et al. 2019). Drought-related mortality may be the result of decreased carbon assimilation and cellular metabolism and increased susceptibility to other stresses, such as disease and insect attack (Bréda et al. 2006; McDowell et al. 2008; Anderegg et al. 2016; McDowell et al. 2022). To respond to water limitation, plants have evolved various strategies to avoid or tolerate stress (Chaves et al. 2003; Verslues et al. 2006; Claeys and Inzé 2013). Plant adaptation to drought stress includes physiological adaptations to maintain water balance throughout dehydration that are controlled by the expression of drought-associated genes. Typical physiological responses include stomatal closure, increased root to shoot growth, reduced leaf expansion, accumulation of solutes, cell wall hardening, and synthesis of protective proteins (Dickson and Tomlinson 1996; Chaves et al. 2003; Verslues et al. 2006). Such responses may increase survival in dry environments, but they may also limit vegetative growth (Verslues et al. 2006; Kaproth and Cavender-Bares 2016). One way to understand complex physiological processes is to use whole-transcriptome sequencing to assay the underlying patterns of gene expression.

Drought tolerance strategies may either be plastic responses induced by water stress or constitutive traits that are continuously expressed regardless of water limitation. Plastic physiological responses can be determined by differentially expressed genes, where the functional roles of up- and down-regulated genes reflect processes that are being turned on or off, respectively. The extent of differential expression can signal the degree of drought tolerance of a species, with fewer differentially expressed genes in drought adapted than drought sensitive species (Madritsch et al. 2019; Mead et al. in review). Other physiological traits may be shaped by high constitutive gene expression throughout treatments with limited plasticity. Such constitutive expression, also described as ‘frontloading’ (Barshis et al. 2013; Rivera et al. 2021), has been observed following stress both within and between populations in rice (Hamann et al. 2024), monkeyflower (Preston et al. 2022), and corals (Barshis et al. 2013). Constitutive expression may facilitate rapid responses.to stress and be favorable in environments where stress events are common (Barshis et al. 2013; Rivera et al. 2021). In the case of water stress, it is possible that dry environments with frequent drought events may put strong selection pressure for high constitutive expression of drought-associated genes and reduced gene expression plasticity. These patterns of gene expression may contribute to drought tolerance strategies and explain differences in drought adaptation among plant species.

Oaks represent an excellent genus with which to pose questions of drought tolerance because they include a range of taxa with different levels of drought adaptation (Kaproth and Cavender-Bares 2016). In general, oak taxa occupy diverse habitats of varying water limitation (Abrams 1990), show high morphological plasticity in drought tolerance (Kaproth and Cavender-Bares 2016), and exhibit variety in water use efficiency between populations within the same species (Roussel et al. 2009). In oaks, several studies have identified gene expression plasticity associated with abiotic stress (Torre et al. 2014; Gugger et al. 2017; Madritsch et al. 2019; Mead et al. 2019). In two prior drought stress studies of *Quercus lobata*, sub-populations showed different patterns of differential gene expression (Gugger et al. 2017; Mead et al. 2019). Overall, Gugger et al. (2017) reported more differentially expressed genes with roughly equivalent numbers of up- and down-regulated genes, while Mead et al. (2019) found slightly more up-than down-regulated genes with putative roles related to water deprivation, heat stimuli, hormone signaling, and gene expression.

Down-regulated genes had photosynthesis- and metabolism-related functions. In general, gene expression studies have largely focused on differential expression in response to drought, which will facilitate plasticity in the affected phenotypes. However, Mead (in review) reported higher levels of constitutive gene expression and reduced plasticity in drought response genes in drought tolerant relative to drought sensitive oak species. Gene expression studies that analyze patterns of up- and down-regulation versus constitutive expression can provide valuable detail on how trees cope with drought stress.

Here, we explore patterns of gene expression in response to experimental drought treatment in an endemic California oak, *Q. douglasii* Hook & Arn (blue oak), a drought tolerant oak tree in California (Huesca et al. 2021). Uniquely, this study of a drought tolerant oak analyzes patterns of differentially expressed genes, and constitutive expression of drought-responsive genes identified from *Q. lobata*, a drought sensitive oak species. We defined patterns of gene expression as constitutive if the initial level of gene expression was greater than zero and remained constant after drought treatment. Specifically, we asked two questions. 1) What are patterns of differential expression of up- and down-regulated genes in response to drought and what are their functional roles? 2) Does *Q. douglasii* additionally show constitutive expression that could mitigate drought stress? We imposed well-watered and dry-down treatments to 24 seedlings germinated from six maternal trees (i.e., six families) from a single natural site in California, and sequenced transcriptomes of all seedlings before and after each treatment (48 libraries). Our study demonstrates that constitutive expression of drought-associated genes is part of the drought-response of *Q. douglasii*, and likely to play a role in other drought-tolerant oaks.

## Materials and Methods

### Study species and sampling site

*Quercus douglasii* Hook & Arn (blue oak) is a deciduous California endemic oak that is widely distributed across the foothills of the Coastal and Eastern Sierra Ranges, and has more coverage and biomass than *Q. lobata* and *Q. agrifolia,* the two other major foothill tree oaks of California (Pavlik 1991). Blue oak is found in drier sites within oak woodland habitats (Pavlik et al. 1991), often occurring on steeper slopes than the latter species (Knops and Koenig 1994). Phylogenetically, *Q. douglasii* is in the section *Quercus* and closely related to the shrub white oaks, especially *Q. john-tuckerii, having emerged* as a species about the same time as the arrival of Mediterranean climate in California (Kim et al. 2018; Hipp et al. 2020). Thus, *Q. douglasii* evolved much later than other sympatric California tree oaks in the same section, including *Quercus*, *Q. lobata* and *Q. garryana* (Hipp et al. 2020, presumably during cooler temperatures and higher summer precipitation. *Q. douglasii* is considered a drought tolerant oak species (Huesca et al. 2021). Overall, blue oak is an ecologically important and biologically useful study system with which to pose questions surrounding drought tolerance.

In October 2013, acorns were sampled from six *Q. douglasii* trees at a single site near O’Neals spanning approximately 1 km^2^ in the Sierra Foothills in California (37.09201, -119.7284; Figure 1). Trees were spaced ∼50m apart in blue oak woodland habitat.

**Figure 1.**
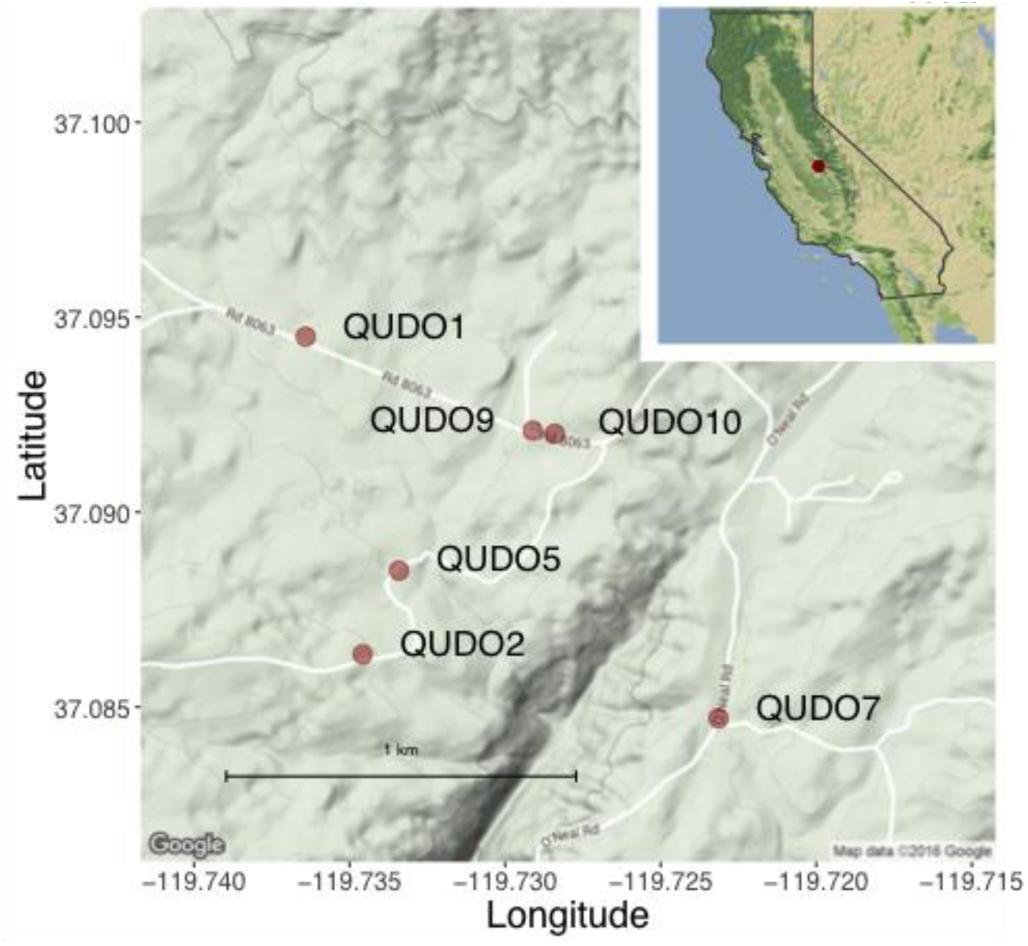
Map of six maternal *Quercus douglasii* source trees whose seedlings provided tissue for the RNA-Seq analysis. Maternal trees were located within approximately 1km^2^ at a single site in the Sierra Nevada Foothills in California.

### Experimental design

Acorns were surface sterilized and planted in the greenhouse at UCLA, with an 18 hr./6 hr. light/dark cycle and 20 – 23°C temperature. Tray positions were randomized every week. After approximately five months in the greenhouse, seedlings were randomly assigned to drought and well-watered treatment groups, with treatment groups and maternal families evenly represented across trays. Well-watered seedlings were watered every few days, while drought seedlings underwent a drought-hardening period for 7 days beginning July 17, 2014, after which they were watered before another dry-down period for 9 days. The goal of the drought-hardening period was to acclimate plants to water stress, such that a second drought period would capture plant response to dehydration beyond initial shock (Vilagrosa et al. 2003; Villar-Salvador et al. 2004). This approach is intended to simulate natural field conditions in which plants are subjected to repeated drought events. Around the treatment period, seedlings were experiencing infection with powdery mildew. Thus, two days prior to the start of the experiment and seven days into the experiment, seedlings were sprayed with a solution (75 mL dish soap, 300 mL ultrapure 98% petroleum oil in 5 gallons water) to remove powdery mildew. Infection appeared to occur indiscriminately, regardless of treatment group.

### RNA extraction and sequencing

Leaf tissue from all seedlings was sampled at two time points for RNA extractions: **Time 1** corresponding to day one at the start of the experiment before soil-drying; and **Time 2** corresponding to day 16 after drought treatment seedlings had undergone two dry-down periods. Leaf tissue was frozen between sheets of dry ice in the greenhouse and transferred to a -80°C freezer.

At each of the two sampling times, twenty-four seedlings from six maternal families were selected for RNA extractions, yielding a total of 48 samples. To reduce variation in seedling growth among samples, we selected seedlings from the middle 70^th^ percentile of the height distribution. To remove polyphenolics and polysaccharides, leaf samples first underwent a lithium chloride/urea-based pre-wash protocol developed for conifers (http://openwetware.org/wiki/Conifer_RNA_prep). The protocol was optimized for *Q. douglasii* samples by implementing the following changes: (1) 50 – 75 mg of tissue was flash-frozen in liquid nitrogen and transferred to frozen 2 mL grinding tubes with two metal beads; (2) tissue was ground twice for 1 min at 30 Hz in flash-frozen adapters; and (3) 1.8 mL Urea/LiCl extraction buffer was added to each tube, allowing recovery of 1.5 mL supernatant. The pre-wash was followed by extraction of total RNA using the RNeasy Plant Mini Kit (QIAGEN, Germantown, MD). An incubation at 56°C with RLT buffer for 2 min, DNase digestion, and an additional 500 μL wash with buffer RPE increased sample purity and yield. Nine samples with low 260/230 ratios on a NanoDrop ND-1000 spectrophotometer (Thermo Fisher Scientific, Waltham, MA) underwent a bead cleanup using an Agencourt AMPure XP kit (protocol B37419AA; Beckman Coulter, Beverly, MA). Total RNA quality and quantity was verified on a 2100 BioAnalyzer using a eukaryotic total RNA Nano Series II assay (Agilent Technologies, Santa Clara, CA) at the UCLA GenoSeq Core.

We used a TruSeq RNA library preparation kit (Illumina, San Diego, CA) to isolate poly-A tail-selected mRNA and convert to cDNA. Twenty-four unique Illumina adapters were used to barcode individual libraries. Libraries were quantified via a Qubit dsDNA BR assay kit (Thermo Fisher Scientific, Waltham, MA), and average fragment size was estimated on a 2100 Agilent Bioanalyzer using a DNA 1000 Series II assay at the UCLA GenoSeq Core. Samples were then normalized and pooled into four libraries, each containing three sets of four samples representing a drought and well-watered seedling at Time 1 and Time 2 for a given maternal family. Libraries underwent single-end 50-bp sequencing on four lanes of an Illumina HiSeq2000 at the Broad Stem Cell Research Center at UCLA.

### Read processing, transcriptome assembly, and annotation

Raw sequence reads were de-multiplexed allowing one mismatch per barcode, followed by several filtering steps. Reads that failed the Illumina quality filter were removed, and remaining reads were converted from qseq to fastq format. Scythe version 0.994 BETA (Buffalo 2011) was used to trim adapter sequence, followed by Sickle version 1.33 (Joshi and Fass 2011) to trim low-quality sequence from read ends falling below an average Phred score of 30 in a sliding window and to remove resulting reads less than 20 bp. Low complexity sequences were removed with the FASTX-Toolkit (Hannon 2010) and processed reads were quality-checked with FastQC (Andrews 2010). To remove ribosomal RNA sequences, reads were mapped to the SILVA rRNA database using BBTools ‘bbmap’ with the parameter ‘outu’ to keep only the unmapped reads. This final quality control step removed 8.3 Gb data. The demultiplexed sequence data have been deposited in the International Nucleotide Sequence Database Collaboration (INSDC) database under the BioProject accession number PRJNA1259526.

A *de novo* transcriptome was assembled using the Trinity platform (Haas et al. 2013). Specifically, we used the genome-guided assembly tool with the *Quercus lobata* reference genome, Valley oak genome 3.0 (Sork et al. 2022) as the closest sequenced species. To functionally annotate the Trinity transcripts and identify candidate coding regions within the transcript sequences, first we used TransDecoder v5.7.1 (Haas,https://github.com/TransDecoder/TransDecoder). Then, we used Trinotate (Bryant et al. 2017) to build an sqlite database with our sequence data and the TransDecoder output, which was used to analyze our sequences against the following databases: BLAST (Altschul et al. 1990), Pfam (Punta et al. 2012), GO (Ashburner et al. 2000), and infernal (Nawrocki and Eddy 2013).

### Differential expression and gene ontology analysis

We performed transcript quantification on gene-level abundance estimates in a genome-free way using Salmon v1.5.2 (Patro et al. 2017) on all read libraries. The relationships within and between biological replicates for each dataset were visually examined using the ‘PtR’ script in the Trinity toolkit (Grabherr et al. 2011; Haas et al. 2013). The transcript count matrix was tested for differential expression using DESeq2 (Love et al. 2014), which uses negative binomial generalized linear models to test for statistical significance. Differential expression was calculated between Time 1 and Time 2 separately for drought and well-watered treatments. We also calculated differential expression using edgeR (Chen et al. 2016), which gave similar results. We used the Benjamini-Hochberg method (Benjamini and Hochberg 1995) to adjust P-values for multiple testing to control the false discovery rate and defined a set of differentially expressed genes with P-values <0.01 and log-fold change >2. To obtain a final set of differentially expressed genes, we first removed any genes present in both drought and well-watered treatments, and then selected only those significant in both DESeq2 and edgeR.

All differentially expressed genes were classified according to their GO (gene ontology) term, creating enriched and depleted gene classes across the treatment groups. Functional enrichment analysis was run using GO-Seq (Young et al. 2010) with GO terms from Trinotate, through the Trinity toolkit ‘analyze_diff_expr.pl’ with the parameters ‘–examine_GO_enrichment –GO_annots – gene_lengths’. This script returns differentially expressed genes for enriched and depleted GO categories for the up- and down-regulated genes in each comparison. Enriched GO terms were further filtered whereby only over-represented terms with a P-value < 0.01 were used for analyses.

To compare patterns of differential expression by gene and GO term, we calculated fold change values (log2FC) for each maternal family using the equation: log2FC = log2(mean_expression_family_1_T2 + 1) – log2(mean_expression_family_1_T1 +1), where the average expression for a gene or GO term was averaged across biological replicates within a family and then averaged by treatment. Second, we tested whether there were differences in expression patterns separately for up- and down-regulated genes between maternal families for individual genes with a repeated measures ANOVA (gene as the repeated measure): expression ∼ treatment * time * family, random effect = gene. When family was significant, we then tested for an effect of maternal family using a generalized linear model and specifying a Tukey-test, by selecting only drought-treatment individuals at Time 2: expression ∼ maternal family + gene. The Benjamini-Hochberg method (Benjamini and Hochberg 1995) was used to adjust P-values for multiple testing to control the false discovery rate.

### Constitutive expression in *Quercus douglasii*

We identified putative drought-responsive genes that responded to drought treatment in *Quercus lobata* seedlings, based on supplementary data files from Gugger et al. (2017) and Mead et al. (2019) (hereafter, drought-responsive genes). Specifically, we found 81 shared protein family (Pfam) categories that were reported to be significantly differentially expressed (p < 0.05 in Gugger et al. (2017) and adjusted P-value < 0.01 in Mead et al. (2019).

To test whether these drought-responsive genes were constitutively expressed in *Q. douglasii*, we measured the expression of *Q. douglasii* genes matching the 81 Pfams across Time 1 and Time 2, separately for well-watered and drought treatments. We first controlled for variance in expression levels by scaling the data (dividing by variance in standard deviation), then tested the significance and magnitude of difference using linear mixed effect models: expression ∼ time, random effect = Pfam. Using a Spearman’s rank correlation, we additionally tested the rank order of Pfam expression, first between Time 1 and Time 2 separately for each treatment and second between well-watered and drought treatments at Time 2. Gene expression is defined as constitutive when the initial level of gene expression was greater than zero and remained more or less constant after drought treatment.

## Data analysis

The following R packages were used in analyses and to make figures, all using the latest versions: tidyverse (Wickham et al. 2019), nlme (Pinheiro and Bates 2025), broom.mixed (Bolker et al. 2024), multcomp (Hothorn et al. 2008), and cowplot (Wilke 2024). This work used computational and storage services associated with the Hoffman2 Cluster, which is operated by the UCLA Office of Advanced Research Computing’s Research Technology Group.

## Results

### Differential expression following drought stress in *Quercus douglasii*

The 48 RNA-seq libraries yielded 914.3 million total raw reads, reducing to 772.9 million reads following quality control. On average, filtered libraries contained 16.1 million ± 2.57 million reads. Among 50,362 total expressed genes, 152 genes were differentially expressed across the drought Time 2 samples in both DESeq2 and edgeR (Figure 2). Most of these differentially expressed genes were down-regulated (123) in the drought-stressed individuals, while a smaller number (29) were up-regulated (Figure 2A). In the well-watered group, essentially zero genes were down-regulated between Times 1 and 2, while a modest number were up-regulated, presumably contributing to the functioning of the seedlings. The log2 fold change values of the significant differentially expressed genes were comparable between down- and up-regulated genes (Figure 2B).

**Figure 2.**
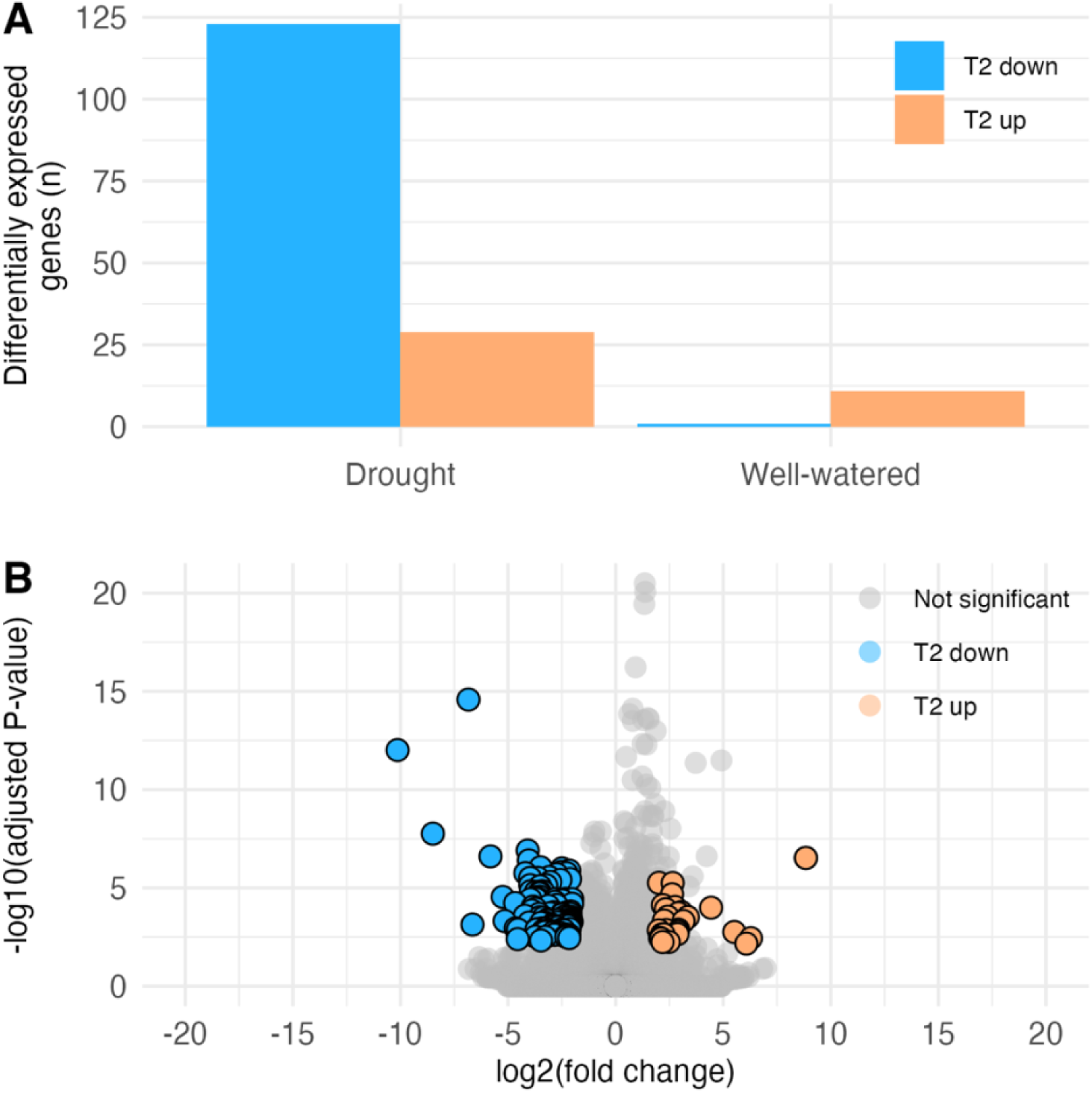
Transcriptional response of *Quercus douglasii* seedlings to experimental drought stress. **A** The number of genes showing differential expression, both up- and down-regulated, in the drought-stressed and the well-watered treatment group at Time 2, as compared with Time 1. Values represent genes which were identified as differentially expressed using negative binomial generalized linear models in both DESeq2 and edgeR, and any genes which were differentially expressed in both drought and well-watered treatments were removed. **B** Volcano plot showing the total differentially expressed genes at Time 2 relative to Time 1 in the drought treatment group. Each point represents a gene’s expression. Orange points are up-regulated genes in response to drought; blue points are down-regulated genes; grey points are genes which are not significantly differentially expressed.

### Significant up-regulation of drought-associated genes

We found 29 genes significantly up-regulated in drought-treated but not well-watered seedlings (Figures 2B, 3). Among these, only one gene ontology (GO) term comprising multiple genes was enriched: four genes were grouped into the enriched molecular function GO term ‘DNA-binding transcription factor activity’. Mostly, up-regulated genes did not match GO terms; instead, we focused upon BLASTp hits. Nine genes returned hits (Table 1), of which many had a putative function in drought response. Specifically, these genes included a cold-regulated protein, monooxygenase, serine acetyltransferase, transcription factor bHLH96, NAC domain-containing protein and transcription factor, reversion to ethylene sensitivity protein, and PXMP2/4 family protein. We did not find an effect of maternal family on expression for up-regulated genes (Table S1).

**Table 1.**
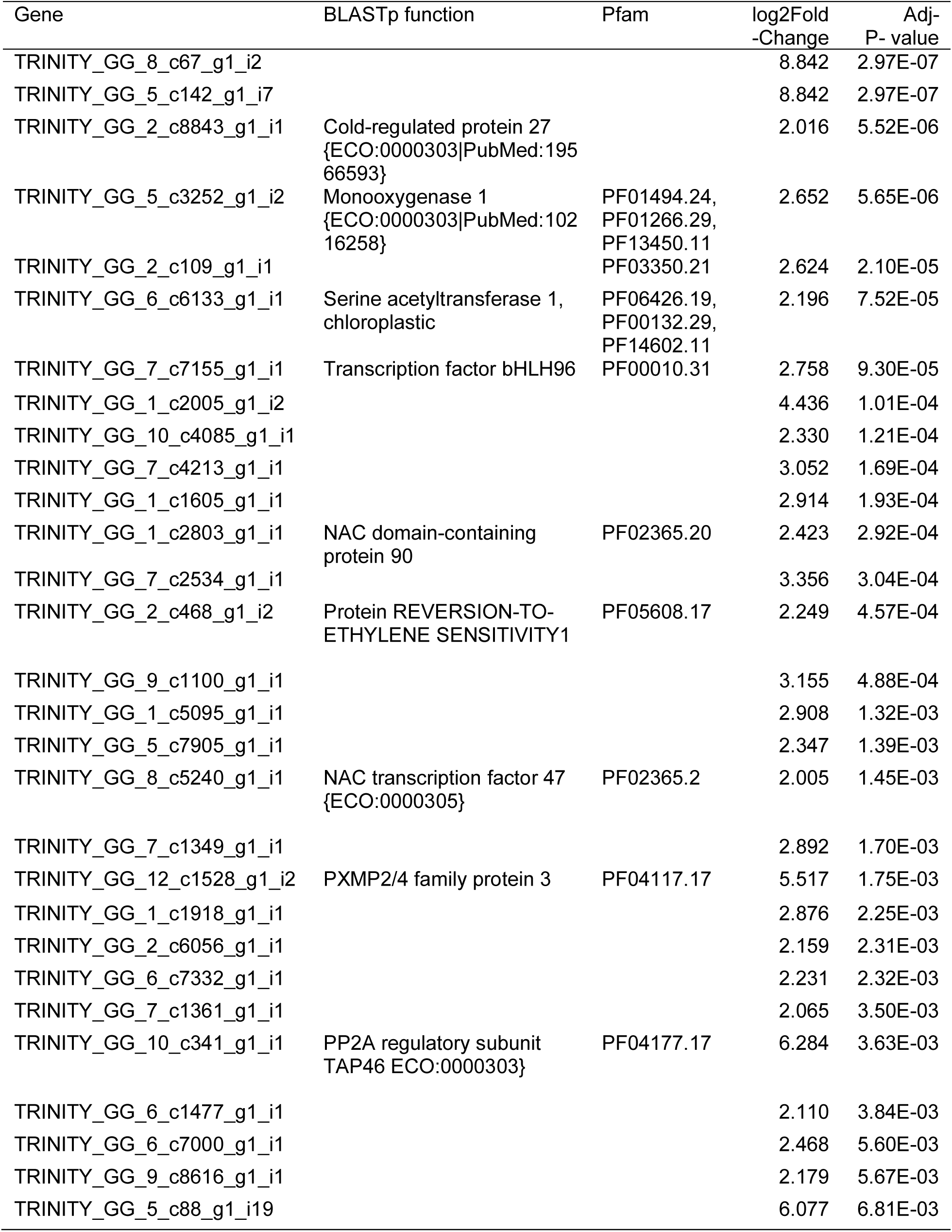
Functional annotations for up-regulated *Quercus douglasii* genes in response to drought.

### Significant down-regulation of genes associated with growth functions

In contrast to the up-regulated genes, we found a significant effect of maternal family on expression of down-regulated genes (Table S2). In addition, conversely to our up-regulated genes, we found three-fold more genes that were down-regulated (123 genes, Figure 2B). in our drought-stressed individuals compared with their well-watered controls. We found a total of 53 enriched GO terms among down-regulated genes (Figure 4 and Table S3). The most strongly down-regulated GO terms largely corresponded to biological process functions, many of which were related to growth, such as ‘lipid catabolic process’, ‘cuticle development’, and various processes related to synthesis of complex sugars found in plant cell walls (Table S3). Enriched GO terms also related to the down-regulation of reproductive processes, including ‘seed coat development’.

**Figure 3.**
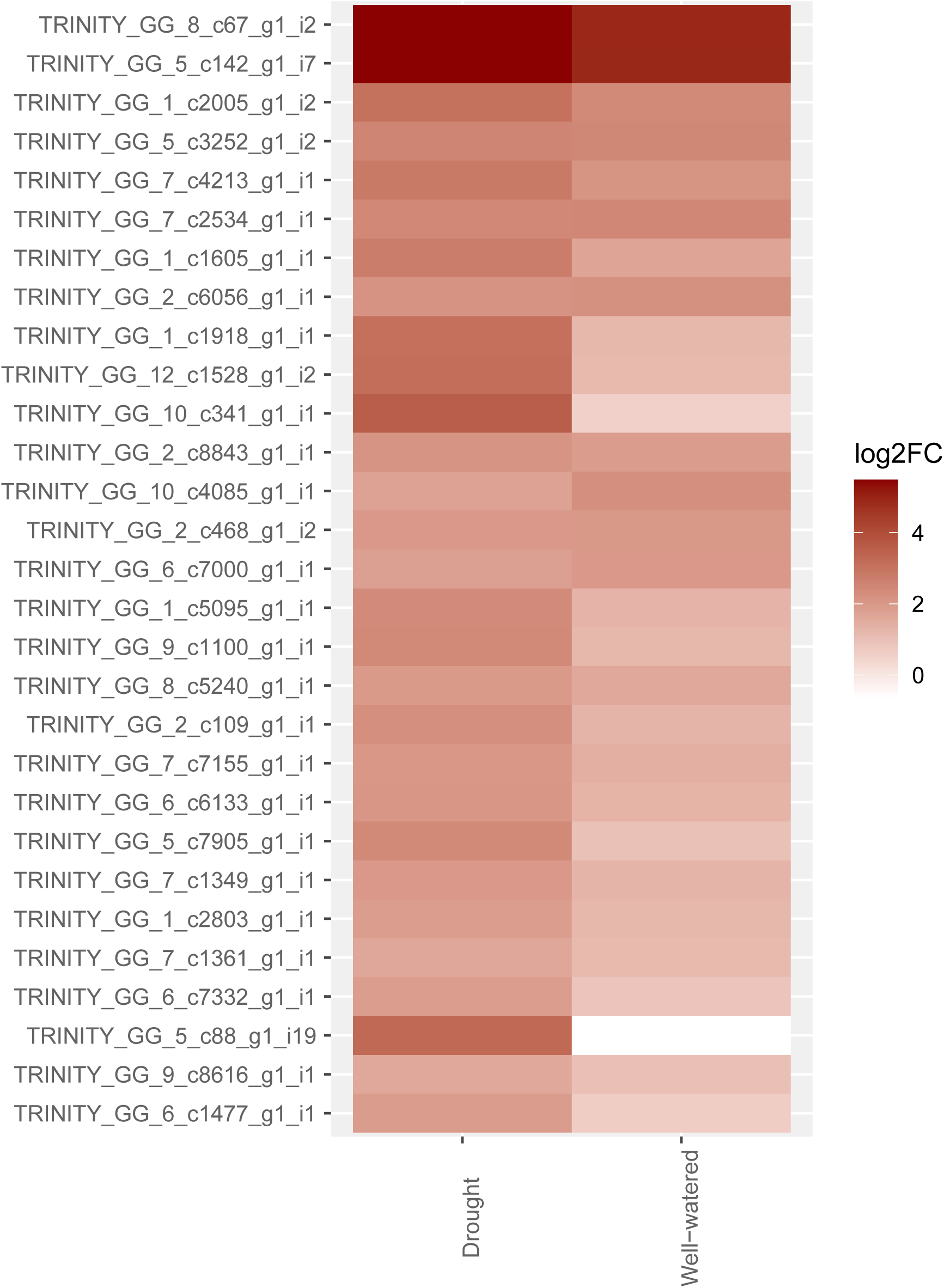
Heatmap of log2 fold change (log2FC) for significantly up-regulated genes following drought treatment. Expression was measured in 24 *Quercus douglasii* seedlings sampled at two timepoints. Fold change was calculated by first taking the average expression for each gene by maternal family, and then calculating the log2 fold change between timepoints for both treatments (See Table 1 for functional annotations of the genes.)

**Figure 4.**
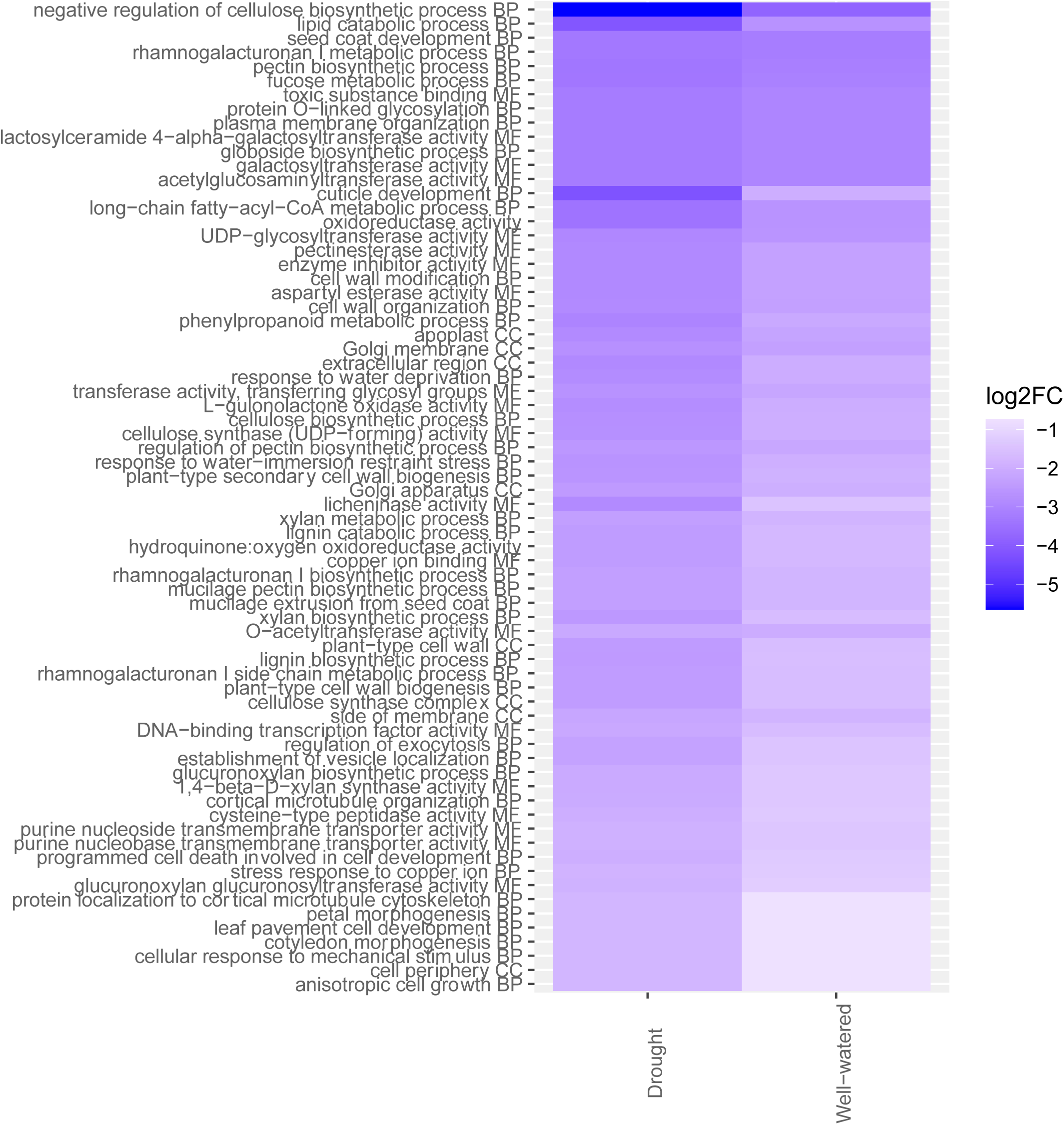
Heatmap of average gene expression for gene ontology (GO) terms enriched within significantly down-regulated genes following drought treatment. Expression was measured in 24 *Quercus douglasii* seedlings sampled at two timepoints. Fold change (log2FC) was calculated by first taking the average expression for each gene for all individuals by maternal family, averaging values for all genes annotated with the given GO term, and finally calculating the log2(fold change) first by family and then between timepoints for both treatments. Abbreviations correspond to GO terms: BP, biological process; CC, cellular component; MF, molecular function.

### Constitutive expression of drought-responsive genes

We compared protein family (Pfam) categories that were differentially expressed across two previous drought treatment studies on a different oak species, *Q. lobata,* to identify important drought genes in *Quercus.* This comparison yielded 81 “drought-responsive” Pfam categories shared by differentially expressed genes spanning both experiments (Table 2). Several Pfam categories with similar functions were repeated in this list, including ABC transporters, glutathione S-transferases, glycogen transferases, KNOX domains, leucine rich repeats, pyridine nucleotide-disulphide oxidoreductases, and tetratricopeptide repeats. Additionally, the drought-responsive Pfams included some stress-responsive functions; for example, a universal stress protein family (PF00582), heat-shock protein 90 (PF00447 and PF02518), and an auxin response factor (PF06507).

**Table 2.**
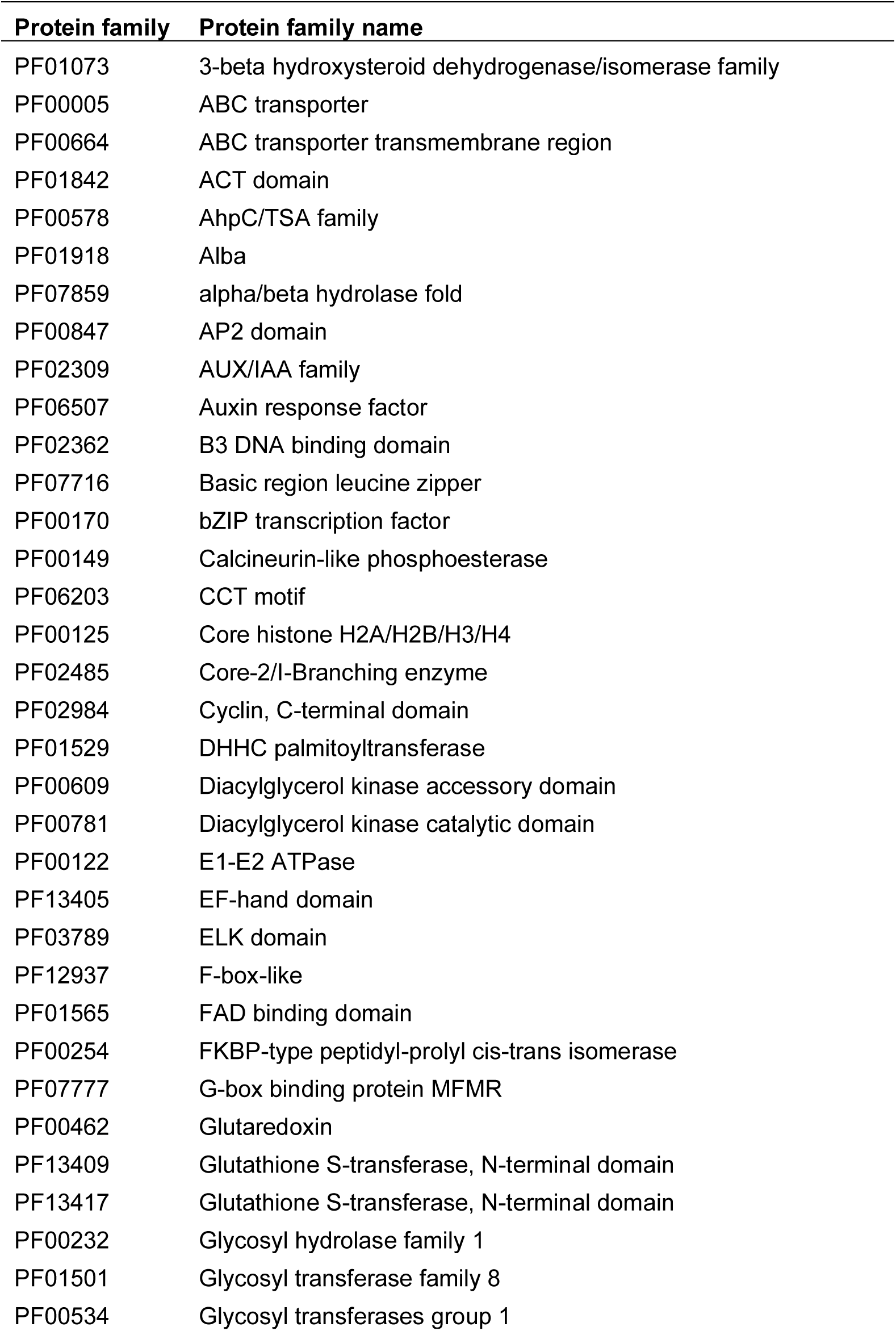
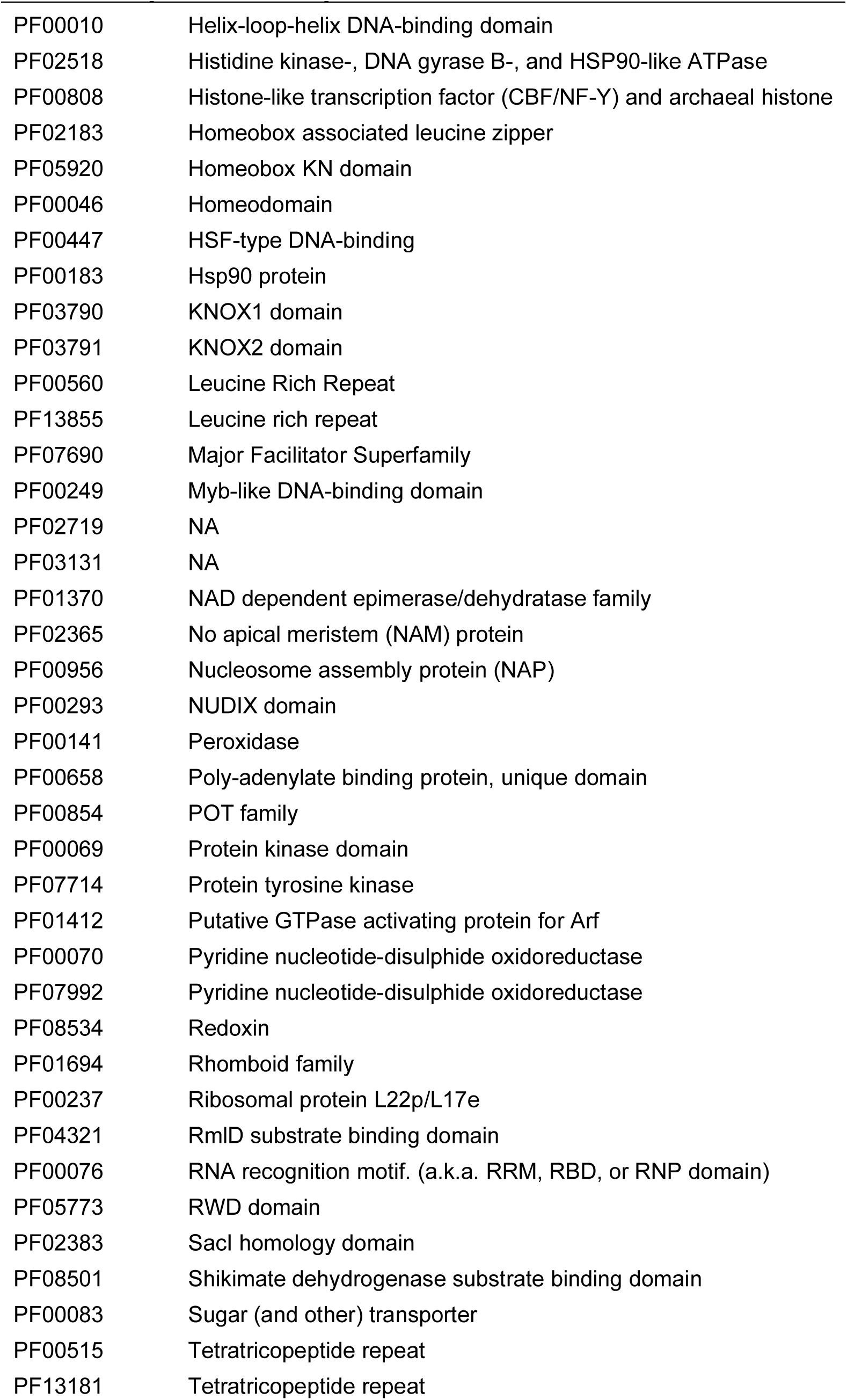
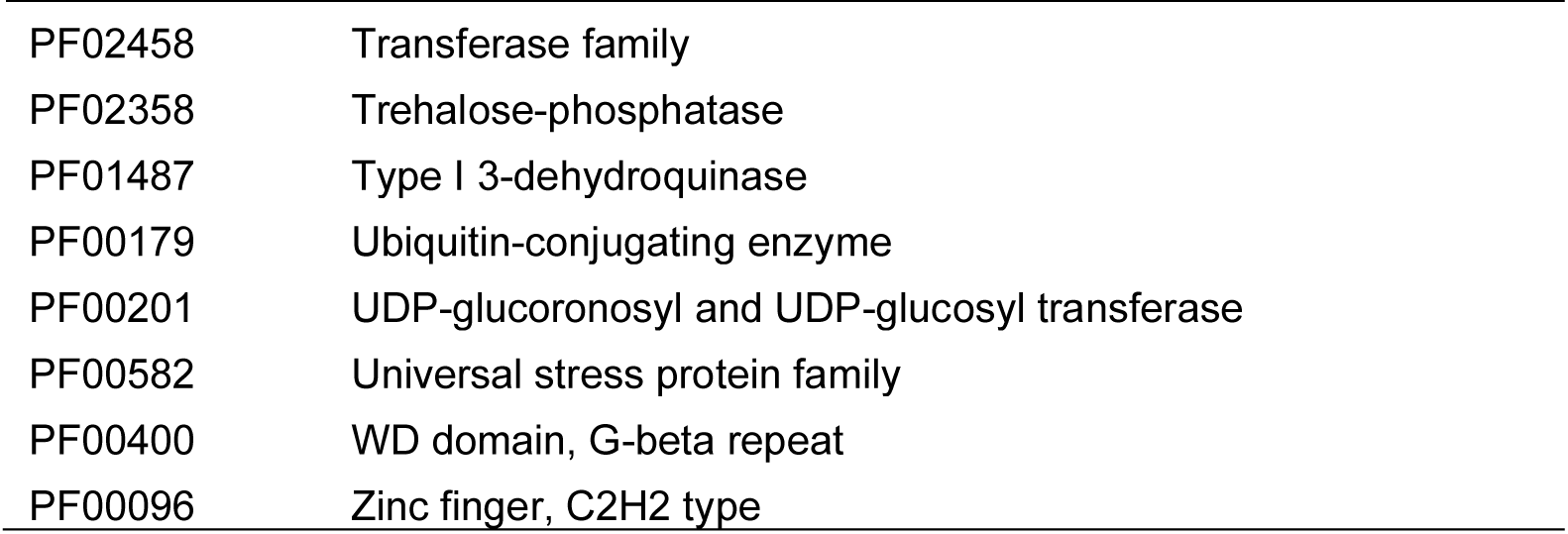
Protein families (Pfams) which were differentially expressed in two drought studies (Gugger et al., 2017, Mead et al., 2019) in *Quercus lobata*, a drought sensitive oak.

| Protein family | Protein family name |
| --- | --- |
| PF01073 | 3-beta hydroxysteroid dehydrogenase/isomerase family |
| PF00005 | ABC transporter |
| PF00664 | ABC transporter transmembrane region |
| PF01842 | ACT domain |
| PF00578 | AhpC/TSA family |
| PF01918 | Alba |
| PF07859 | alpha/beta hydrolase fold |
| PF00847 | AP2 domain |
| PF02309 | AUX/IAA family |
| PF06507 | Auxin response factor |
| PF02362 | B3 DNA binding domain |
| PF07716 | Basic region leucine zipper |
| PF00170 | bZIP transcription factor |
| PF00149 | Calcineurin-like phosphoesterase |
| PF06203 | CCT motif |
| PF00125 | Core histone H2A/H2B/H3/H4 |
| PF02485 | Core-2/l-Branching enzyme |
| PF02984 | Cyclin, C-terminal domain |
| PF01529 | DHHC palmitoyltransferase |
| PF00609 | Diacylglycerol kinase accessory domain |
| PF00781 | Diacylglycerol kinase catalytic domain |
| PF00122 | E1-E2 ATPase |
| PF13405 | EF-hand domain |
| PF03789 | ELK domain |
| PF12937 | F-box-like |
| PF01565 | FAD binding domain |
| PF00254 | FKBP-type peptidyl-prolyl cis-trans isomerase |
| PF07777 | G-box binding protein MFMR |
| PF00462 | Glutaredoxin |
| PF13409 | Glutathione S-transferase, N-terminal domain |
| PF13417 | Glutathione S-transferase, N-terminal domain |
| PF00232 | Glycosyl hydrolase family 1 |
| PF01501 | Glycosyl transferase family 8 |
| PF00534 | Glycosyl transferases group 1 |

| **Protein family** | **Protein family name** |
| --- | --- |
| PF00010 | Helix-loop-helix DNA-binding domain |
| PF02518 | Histidine kinase-, DNA gyrase B-, and HSP90-like ATPase |
| PF00808 | Histone-like transcription factor (CBF/NF-Y) and archaeal histone |
| PF02183 | Homeobox associated leucine zipper |
| PF05920 | Homeobox KN domain |
| PF00046 | Homeodomain |
| PF00447 | HSF-type DNA-binding |
| PF00183 | Hsp90 protein |
| PF03790 | KNOX1 domain |
| PF03791 | KNOX2 domain |
| PF00560 | Leucine Rich Repeat |
| PF13855 | Leucine rich repeat |
| PF07690 | Major Facilitator Superfamily |
| PF00249 | Myb-like DNA-binding domain |
| PF02719 | NA |
| PF03131 | NA |
| PF01370 | NAD dependent epimerase/dehydratase family |
| PF02365 | No apical meristem (NAM) protein |
| PF00956 | Nucleosome assembly protein (NAP) |
| PF00293 | NUDIX domain |
| PF00141 | Peroxidase |
| PF00658 | Poly-adenylate binding protein, unique domain |
| PF00854 | POT family |
| PF00069 | Protein kinase domain |
| PF07714 | Protein tyrosine kinase |
| PF01412 | Putative GTPase activating protein for Arf |
| PF00070 | Pyridine nucleotide-disulphide oxidoreductase |
| PF07992 | Pyridine nucleotide-disulphide oxidoreductase |
| PF08534 | Redoxin |
| PF01694 | Rhomboid family |
| PF00237 | Ribosomal protein L22p/L17e |
| PF04321 | RmID substrate binding domain |
| PF00076 | RNA recognition motif. (a.k.a. RRM, RBD, or RNP domain) |
| PF05773 | RWD domain |
| PF02383 | SacI homology domain |
| PF08501 | Shikimate dehydrogenase substrate binding domain |
| PF00083 | Sugar (and other) transporter |
| PF00515 | Tetratricopeptide repeat |
| PF13181 | Tetratricopeptide repeat |
| PF02458 | Transferase family |
| PF02358 | Trehalose-phosphatase |
| PF01487 | Type I 3-dehydroquinase |
| PF00179 | Ubiquitin-conjugating enzyme |
| PF00201 | UDP-glucuronosyl and UDP-glucosyl transferase |
| PF00582 | Universal stress protein family |
| PF00400 | WD domain, G-beta repeat |
| PF00096 | Zinc finger, C2H2 type |

To determine whether drought-responsive genes in *Q. lobata* were also drought-responsive in *Q. douglasii*, we retrieved transcripts matching these 81 Pfams and analysed their mean expression at Time 1 and 2 in both well-watered and drought treatments. Unlike in *Q. lobata,* we saw no difference in mean expression of these drought-responsive Pfams in either treatment (Figure 5, p > 0.05), suggesting their differential expression is a *Q. lobata*-specific, not *Q. douglasii*, drought stress response. Expression levels of each Pfam were similar both between Time 1 and Time 2 within each treatment, and between Time 2 in drought and well-watered treatments (Spearman’s rank correlation rho: drought Time 1 vs Time 2, rho = 0.96, p < 2.2e-16; well-watered Time 1 vs Time 2, rho = 0.95, p < 2.2e-16; Time 2 drought vs well-watered, rho = 0.99, p < 2.2e-16).

**Figure 5.**
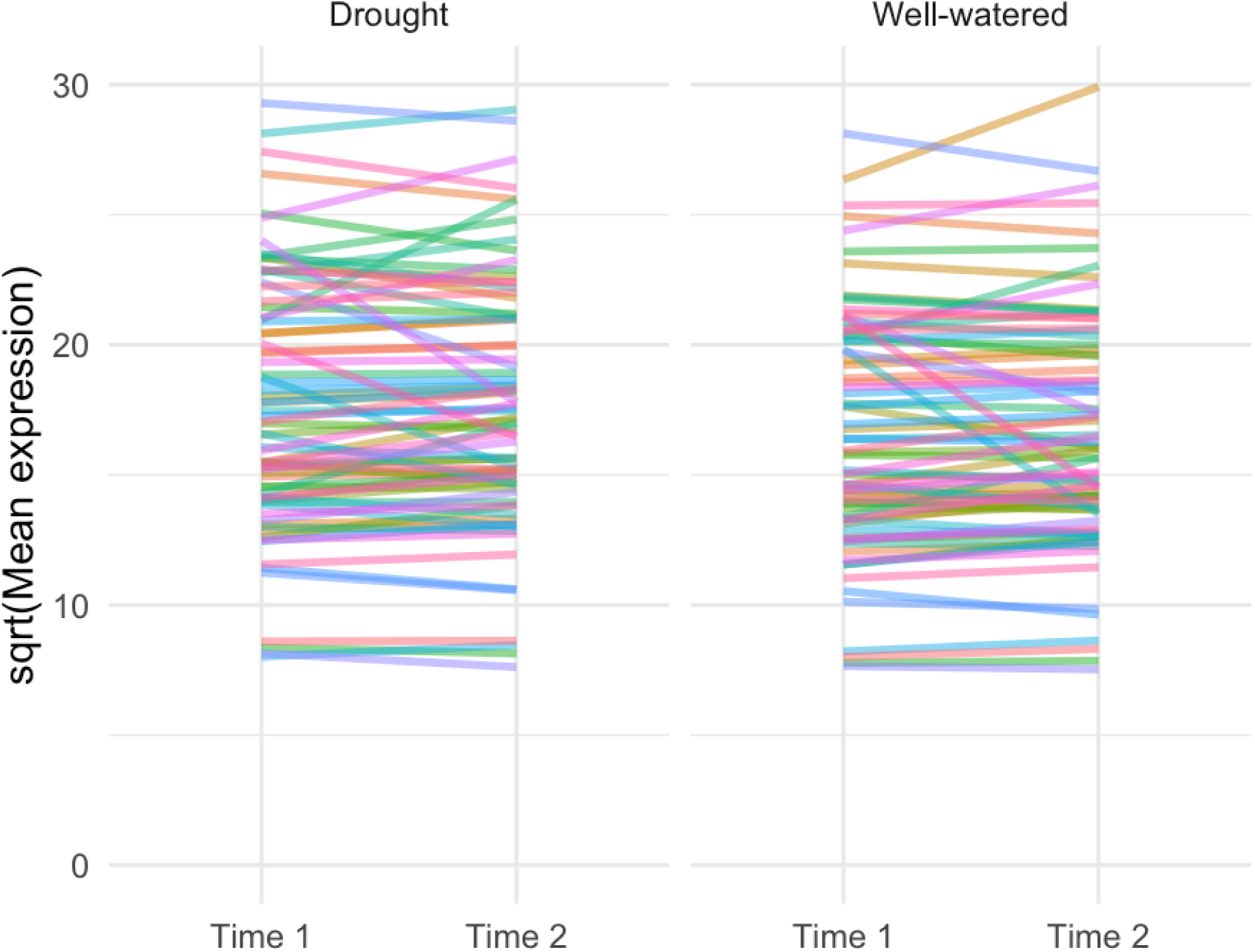
Gene expression for 81 drought-responsive protein families (Pfams) in *Quercus douglasii* drought and well-watered treatments. Lines represent mean expression of 25,630 transcripts (calculated by negative binomial generalized linear models in DESeq2) across Time 1 and Time 2 for genes within each Pfam category; line shading indicates Pfam. Neither drought nor well-watered show an effect between time points (drought: slope = 0.007, P > 0.05; well-watered: slope = 0.01, P > 0.05).

## Discussion

Our study illustrates an intriguing response to drought stress in seedlings collected from a California population of a drought tolerant oak, *Q. douglasii*. We found more down-regulated genes than up-regulated genes, and many drought-responsive genes that appear to be constitutively expressed. Down-regulated genes were typically involved in growth, indicating that drought tolerance may come at the cost of growth. Few up-regulated genes may be explained by the high number of drought-responsive genes that were already expressed at the start of the experiment. These genes comprise 81 drought-responsive protein families (Pfams) that were identified in *Q. lobata*. Results from our drought stress experiment highlight that *Q. douglasii* seedlings respond to drought both through gene expression plasticity triggered by low soil moisture and through genes with high constitutive expression. Thus, in a drought-adapted species exposed to constant water stress, constitutive gene expression may be a key aspect of their adaptation to low water environments.

### Gene expression plasticity in *Quercus douglasii*

The seedlings in this study showed few differentially expressed genes in response to drought, and with the majority down-regulated. This finding of an overall low number of differentially expressed genes could relate to the fact that *Q. douglasii* is a drought-tolerant species. Similar to our study, two independent studies in oaks in Europe and California (Madritsch et al. 2019; Mead et al. in review) found that drought tolerant oak species (*Q. ilex* and *Q. pubescens* in Europe; *Q. palmeri, Q. chrysolepis, Q. durata* and *Q. agrifolia* in California exhibited fewer differentially expressed genes than less drought tolerant species (*Q. robur* in Europe; *Q. lobata* and *Q. kelloggii* in California) in response to soil dry down treatment. We can gain some insight about why drought-tolerant species have fewer differentially expressed genes from the European study (Madritsch et al. 2019), who conclude that the two drought-tolerant Mediterranean species exhibited drought avoidance while *Q. robur* encountered severe stress, as evidenced by lowered root growth. Thus, it seems that drought tolerant oaks may not need to respond to drought stress by increasing gene expression.

Gene functions for up-regulated genes were similar to those reported previously for oaks (Gugger et al. 2017; Madritsch et al. 2019; Mead et al. 2019), with most gene functions broadly associated with drought response. For example, serine acetyltransferase 1 was up-regulated, an important component of the cysteine synthase complex which in turn regulates the biosynthesis of abscisic acid (ABA), a major hormone known to mediate drought response. Various studies have reported the up-regulation of serine acetyltransferases in response to abiotic stress (Kurt et al. 2021; Wang et al. 2024). Additionally, the ethylene-signaling protein RTE1 (REVERSION-TO-ETHYLENE SENSITIVITY 1) was up-regulated, which reduces sensitivity to ethylene (another stress hormone) and improves drought tolerance when overexpressed in Arabidopsis and maize (Shi et al. 2015).

Finally, two transcription factors were up-regulated, a basic helix-loop-helix (bHLH) and an NAC transcription factor, which both have well-reported roles in plant response to abiotic stress, e.g., bHLH promotes drought tolerance in tomato by switching on genes encoding antioxidants, ABA-signaling molecules, and stress-related proteins (Liang et al. 2022) and NAC regulates endogenous ABA in Arabidopsis (Jensen et al. 2013). The preponderance of differential expression of drought-related genes indicates that soil-drying triggered genes that respond to drought stress.

The fact that in our study most differentially expressed genes were down-regulated is also reported in other species, such as two desert species (Long et al. 2014; Ma et al. 2015) and willows (Pucholt et al. 2015). In terms of down-regulated gene functions, our findings were consistent with other studies that reported an inhibition of photosynthesis, carbohydrate metabolism, and cell division under drought stress (Hayano-Kanashiro et al. 2009; Gugger et al. 2017; Zhang et al. 2018). Carbohydrate metabolism and biosynthesis are critical plant processes for capturing energy produced during photosynthesis, and its substrates are known to play a role in responses to drought stress as well as providing energy. Carbohydrate levels in tissues can be changed by differential expression of the genes underlying carbohydrate metabolism and biosynthesis. We report the down-regulation of five GO terms related to carbohydrate metabolism: rhamnogalacturonan I side chain metabolic process; long-chain fatty-acyl-CoA metabolic process; xylan metabolic process; fucose metabolic process; rhamnogalacturonan I metabolic process. We additionally report the down-regulation of ten GO terms related to carbohydrate biosynthetic processes: xylan biosynthetic process; lignin biosynthetic process; negative regulation of cellulose biosynthetic process; cellulose biosynthetic process; glucuronoxylan biosynthetic process; regulation of pectin biosynthetic process; pectin biosynthetic process; rhamnogalacturonan I biosynthetic process; mucilage pectin biosynthetic process; globoside biosynthetic process. Therefore, carbohydrate activities were significantly repressed under water restriction, which would result in a carbon deficiency. This response in turn would affect the chloroplast and cell wall and other important developmental functions. We found a preponderance of growth or metabolism processes were being shut down with water stress in *Q. douglasii,* which could reflect a trade-off between growth and drought tolerance mechanisms.

Within our sample of seedlings from one population, we discovered that the down-regulated genes exhibited variable gene expression in response to drought (treatment × time × maternal family interaction), while the up-regulated genes did not. This finding illustrates variation within our population for gene expression plasticity, namely that we see no difference between maternal family in the up-regulation of a small number of drought-responsive genes while differences in down-regulated genes involved in growth could indicate differences in stress tolerance between maternal families, which selection could act on in the future. Rivera et al. (2021) define tolerance as survival or maintained organismal functioning under stress, and the magnitude of this down-regulation affecting how well each *Q. douglasii* family can maintain growth could be a factor contributing to tolerance within each family. The effectiveness of the drought response may depend on how plants perceive drought stress signals and respond via changes in gene expression.

### Constitutive gene expression as a drought response strategy

The finding that many drought-responsive genes in *Q. douglasii* showed high baseline and not differential expression was unexpected, and we were intrigued by the hypothesis of constitutive expression as a drought-response strategy. Therefore, we designed an additional test for higher constitutive expression of the drought-responsive Pfams in *Q. douglasii* as compared with the less drought-adapted *Q. lobata.* With a transcriptome that we generated using *Q. lobata* sequence data from select sites studied by Mead et al. (2019), we calculated differential expression between drought treatments (see supplementary methods). We selected sites with both similar (Centerville) and contrasting (Malibu Creek) climates to that of *Q. douglasii* (O’Neals, Figure 6A). We filtered genes that were differentially expressed in *Q. lobata* to only include those matching our 20 drought-responsive Pfams (80 *Q. lobata* genes). We found that drought treatment did not change the expression of these drought-responsive Pfams in either *Q. douglasii* from O’Neals nor *Q. lobata* from Centerville, but did increase their expression in *Q. lobata* from Malibu Creek (Figure 6B-C).

**Figure 6.**
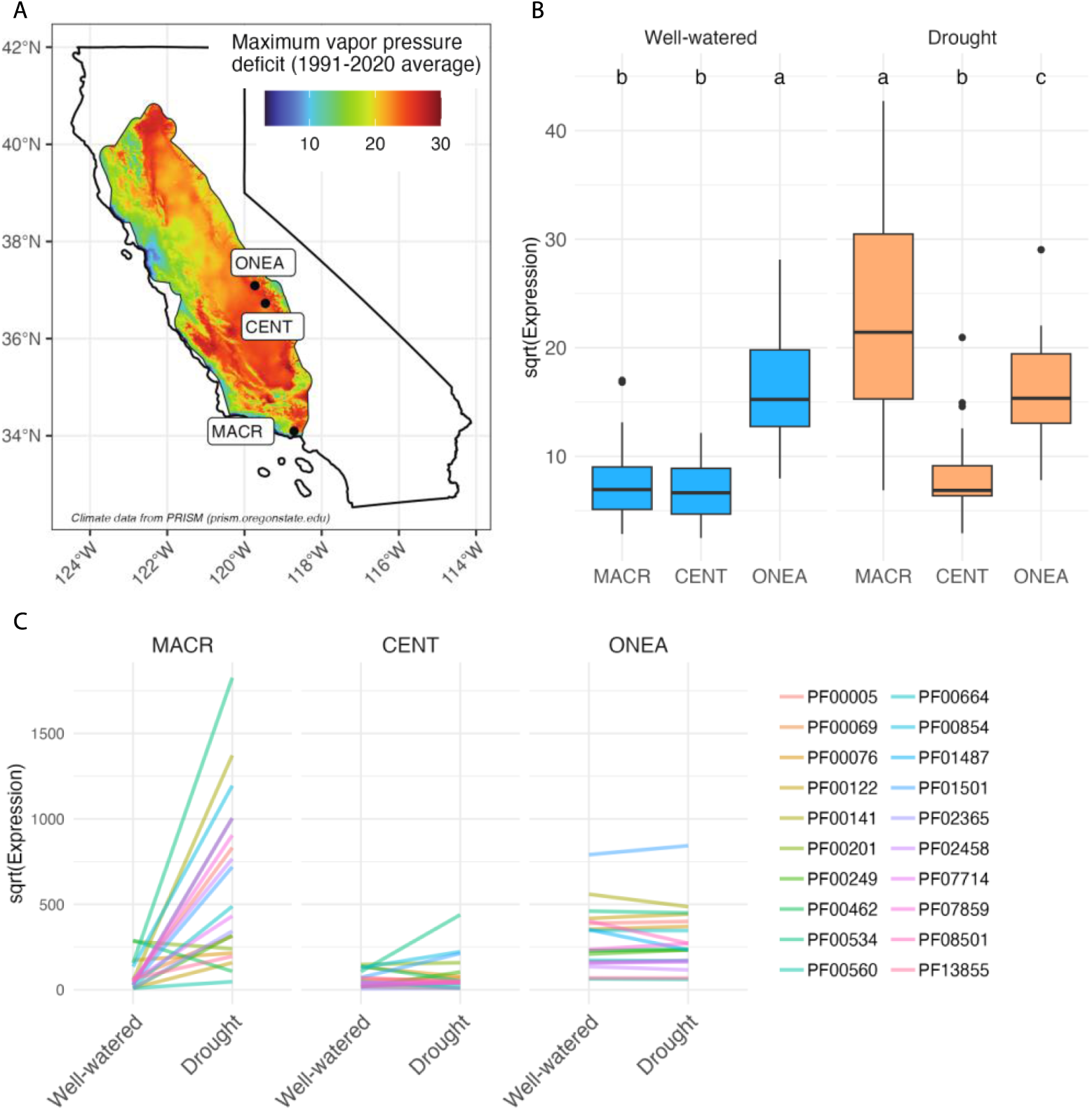
Gene expression for a subset of drought-responsive protein families (Pfams) in *Quercus lobata* and *Q. douglasii* drought and well-watered treatments. **(A)** Map of sites where acorns were sampled in California overlayed with maximum vapor pressure deficit (the difference between the amount of moisture in the air and how much moisture the air can hold at saturation, where higher values indicate drier sites) in the *Q. lobata* and *Q. douglasii* species range (1991-2020 average). Climate data is from PRISM (prism.oregonstate.edu). *Q. lobata* acorns were collected from Malibu Creek (MACR) and Centerville (CENT) and *Q. douglasii* acorns were collected from O’Neals (ONEA). Vapor pressure deficit is lower in Malibu Creek as compared with Centerville and O’Neals. **(B)** Mean expression under well-watered and drought treatments of 20 drought-responsive Pfams that were differentially expressed in the first two sites for *Q. lobata* (MACR and/ or CENT) and present in the third site in figure for *Q. douglasii* (ONEA). Mean expression was calculated by negative binomial generalized linear models in DESeq2 across well-watered and drought treatments for genes within each Pfam category. The 20 drought-responsive Pfams were those comprising >1 differentially expressed *Q. lobata* gene. Letters represent significant differences between sites across each treatment (post-hoc Tukey test with Bonferroni adjusted P-values: expression * site + (1+Pfam)). (**C)** Mean expression of 20 drought-responsive Pfams under well-watered and drought treatments by site. Lines represent mean expression across Time 1 and Time 2 for genes within each Pfam category; line shading indicates Pfam. The 20 Pfams correspond to 80 differentially expressed genes in *Q. lobata* and 2382 genes (and 3731 transcripts) in *Q. douglasii.* Mean expression of *Q. lobata* individuals at Malibu Creek (MACR) show an effect in response to drought (slope = 0.84, p < 0.001) whilst we see no effect for *Q. lobata* individuals at Centerville (slope = 0.17, p > 0.05) nor *Q. douglasii* individuals at O’Neals (slope = 0.01, p > 0.05). The slopes significantly differ by both site and by Pfam (ANOVA: site, p < 0.001; Pfam, p < 0.01).

Interestingly, these Pfams had significantly higher constitutive expression under both the well-watered and drought treatments in *Q. douglasii* compared with *Q. lobata* at both sites (Figure 6B). This suggests that the lack of a transcriptional response to drought treatment in *Q. douglasii* (Figure 5) was because the more drought-tolerant *Q. douglasii* demonstrates high constitutive expression, or front-loading, of drought-responsive genes compared with the less drought-tolerant *Q. lobata.* This strategy could at least partly explain the species’ drought tolerant nature, and reinforces the value of analyzing constitutive expression in individuals sampled from multiple populations.

We conclude that constitutive gene expression in *Q. douglasii* might be an adaptation to limited water availability. This interpretation is supported by a separate gene expression study across six California oak species ranging in drought tolerance (Mead et al. in review), in which higher constitutive expression was reported for a subset of genes in drought tolerant oak species. Together, these findings highlight that drought response could result from plasticity, or local adaptation, or both, in terms of gene expression. Gene expression links genotype to adaptive phenotypes, making gene expression plasticity an important functional response to environmental change across generations (de Nadal et al. 2011). However, another rarely explored yet potentially important component of functional response to drought could be keeping genes turned on.

## Conclusion

Our study provides evidence that *Q. douglasii* demonstrates higher constitutive expression of drought-responsive genes compared with the drought sensitive *Q. lobata.* In a drought tolerant species, water limitation could be less stressful thanks to frontloading, in which higher constitutive expression of specific genes promotes tolerance to stress by maintaining homeostasis and cellular integrity. In contrast, a drought sensitive species may show lower baseline expression of drought-responsive genes because it may experience this type of stress less frequently, resulting in a strong gene expression plasticity response when water limitation is encountered. Our study identified drought-responsive protein families that underlie drought stress response in both *Q. douglasii* and *Q. lobata,* albeit via different patterns of gene expression. To our knowledge, only Mead et al. (in review) and this study have identified constitutive gene expression as a potential adaptation for tolerance to drought stress in any tree species, illustrating that plasticity in gene expression is not the only response strategy that will allow trees to tolerate drought stress.

## Data availability

All raw sequence data have been deposited in the relevant International Nucleotide Sequence Database Collaboration (INSDC) database with the BioProject ID PRJNA1259526. The annotated *Q. douglasii* transcriptome analyzed in this manuscript is published in a publicly available Zenodo digital repository (https://github.com/lilypeck/qdoug-drought-expression/TBD), along with source data, differential expression output files, custom analysis and plotting scripts. This study also re-analyzed publicly available data from Mead et al. (2019).

## Acknowledgments

We acknowledge the California Indigenous groups (past, present, future) who live in relationship to the oak ecosystems that we study. We thank Drs. Lynn Sweet and Frank Davis for assistance with blue oak acorn samples and Sorel Fitz-Gibbon for bioinformatic advice. We thank Kristina Beckley for lab and greenhouse assistance.

## Author contributions (CRediT taxonomy)

Conceptualization: Stephanie Steele and Victoria L.

Sork Methodology: Stephanie Steele

Investigation: Stephanie Steele

Formal analysis: Lily D. Peck and Stephanie Steele

Data curation: Lily D. Peck

Visualization: Lily D. Peck

Writing – original draft: Stephanie Steele and Lily D. Peck, VL Sork

Writing – review and editing: Stephanie Steele, Lily D. Peck and Victoria L. Sork

Funding acquisition: VL Sork, S Steele

## Supplementary Material

### Supplemental Methods

#### Differential expression in *Quercus lobata*

We downloaded the raw sequencing reads from the Mead et al. (2019) data repository (https://doi.org/10.5068/D1HH31) for two *Q. lobata* populations: Centerville, CENT and Malibu Creek, MACR. We performed transcript quantification and differential expression analyses exactly as we did for *Q. douglasii.* To ensure consistency in our methods, we analyzed differential expression separately for each *Q. lobata* site and combined the lists of differentially expressed genes. We selected only differentially expressed genes that matched our drought-responsive Pfams, and compared their expression levels to genes comprising the same Pfams in *Q. douglasii*.

We first controlled for variance in expression levels by scaling the data, then tested the significance of overall expression differences between treatments by fitting an ANOVA: expression ∼ site * treatment, random effect = Pfam. The interaction was significant and so we tested for an effect of site separately for each treatment using a generalized linear model and specifying a Tukey-test: expression ∼ site, random effect = Pfam. The Bonferroni method was used to adjust P-values for multiple testing to control the false discovery rate. To test the significance and magnitude of expression differences by Pfam we used linear mixed effect models: expression ∼ treatment, random effect = Pfam.

## Supplementary Tables

**Table S1.**
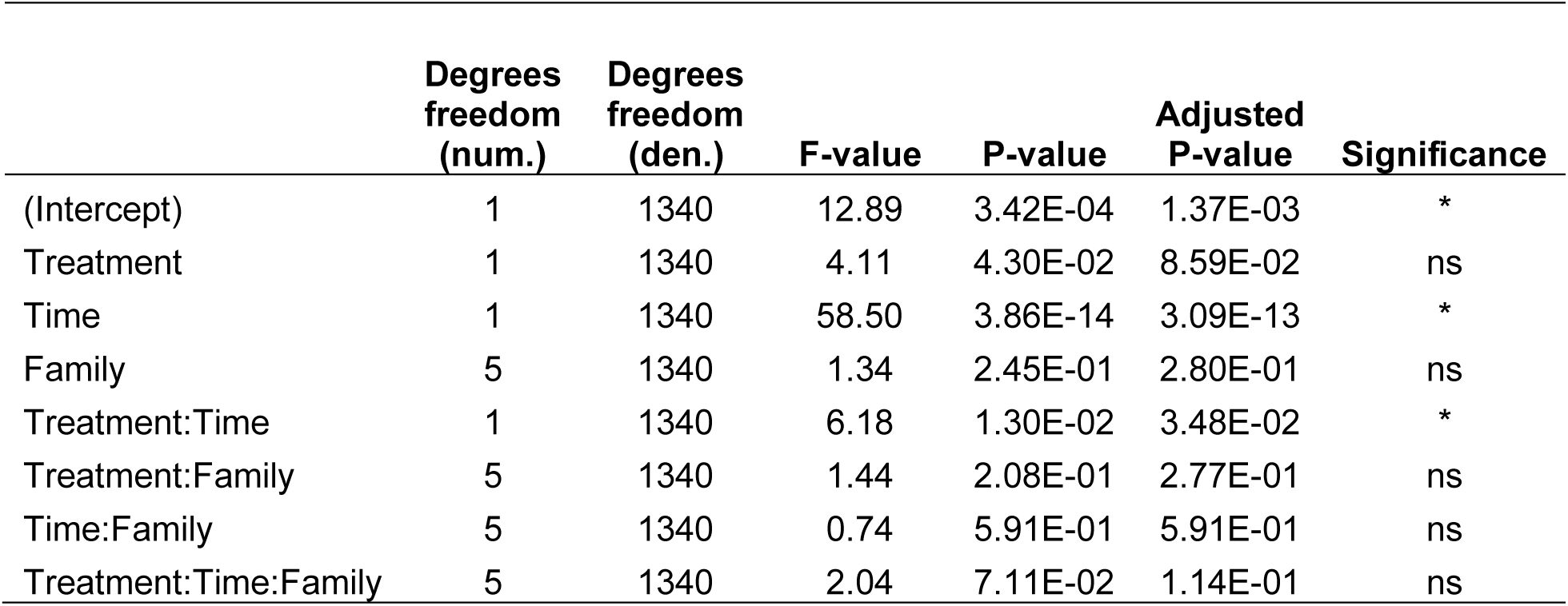
Results of ANOVA, using up-regulated genes, comparing an effect of treatment, time, and maternal family of *Q. douglasii* seedlings in gene expression response to drought. Model: anova(lme(Expression ∼ Treatment * time * Family, random effect = gene)). Abbreviations: num., numerator; den., denominator.

**Table S2.** (**A)**. Results of ANOVA, using down-regulated genes, comparing an effect of treatment, time, and maternal family of *Q. douglasii* seedlings in gene expression response to drought. Model: anova(lme(Expression ∼ Treatment * time * Family, random effect = gene)). Abbreviations: num., numerator; den., denominator. (**B)**. Results of a post-hoc Tukey test comparing pairwise maternal family effects. Model: aov(Expression ∼ Family + (1+gene)). The maternal families are QUDO_1, QUDO_2, QUDO_5, QUDO_7, QUDO_9 and QUDO_10.

| **A** | Degrees<br>freedom<br>(num.) | Degrees<br>freedom<br>(den.) | F-value | P-value | Adjusted<br>P-value | Signifi-<br>cance |
| --- | --- | --- | --- | --- | --- | --- |
| (Intercept) | 1 | 5758 | 54.91 | 1.44E-13 | 2.89E-13 | * |
| Treatment | 1 | 5758 | 1.17 | 2.79E-01 | 2.79E-01 | ns |
| time | 1 | 5758 | 373.98 | 0.00E+00 | 0.00E+00 | * |
| Family | 5 | 5758 | 24.59 | 0.00E+00 | 0.00E+00 | * |
| Treatment:time | 1 | 5758 | 20.95 | 4.81E-06 | 5.50E-06 | * |
| Treatment:Family | 5 | 5758 | 27.79 | 0.00E+00 | 0.00E+00 | * |
| time:Family | 5 | 5758 | 6.61 | 3.83E-06 | 5.10E-06 | * |
| Treatment:time:Family | 5 | 5758 | 8.85 | 2.23E-08 | 3.57E-08 | * |

| **B. Pairwise contrast between families.** |  |  |  |  |  |  |  |
| --- | --- | --- | --- | --- | --- | --- | --- |
| Mat. Fam.<br>1 | Mat. Fam.<br>2 | null.<br>value | Esti-<br>mate | Std. error | T-statistic | Adjusted<br>P-value | Signifi-<br>cance |
| QUDO_10 | QUDO_1 | 0 | -10.5 | 4.52 | -2.32 | 0.309 |  |
| QUDO_2 | QUDO_1 | 0 | -14.4 | 4.52 | -3.18 | 0.0229 | * |
| QUDO_5 | QUDO_1 | 0 | 5.63 | 4.52 | 1.24 | 1 |  |
| QUDO_7 | QUDO_1 | 0 | -16.9 | 4.52 | -3.74 | 0.00292 | ** |
| QUDO_9 | QUDO_1 | 0 | -15.9 | 4.52 | -3.51 | 0.00696 | ** |
| QUDO_2 | QUDO_10 | 0 | -3.88 | 4.52 | -0.858 | 1 |  |
| QUDO_5 | QUDO_10 | 0 | 16.1 | 4.52 | 3.56 | 0.00572 | ** |
| QUDO_7 | QUDO_10 | 0 | -6.41 | 4.52 | -1.42 | 1 |  |
| QUDO_9 | QUDO_10 | 0 | -5.39 | 4.52 | -1.19 | 1 |  |
| QUDO_5 | QUDO_2 | 0 | 20 | 4.52 | 4.42 | 0.000161 | *** |
| QUDO_7 | QUDO_2 | 0 | -2.53 | 4.52 | -0.56 | 1 |  |
| QUDO_9 | QUDO_2 | 0 | -1.51 | 4.52 | -0.333 | 1 |  |
| QUDO_7 | QUDO_5 | 0 | -22.5 | 4.52 | -4.98 | 0.0000108 | *** |
| QUDO_9 | QUDO_5 | 0 | -21.5 | 4.52 | -4.75 | 0.0000334 | *** |
| QUDO_9 | QUDO_7 | 0 | 1.02 | 4.52 | 0.226 | 1 |  |

**Table S3.**
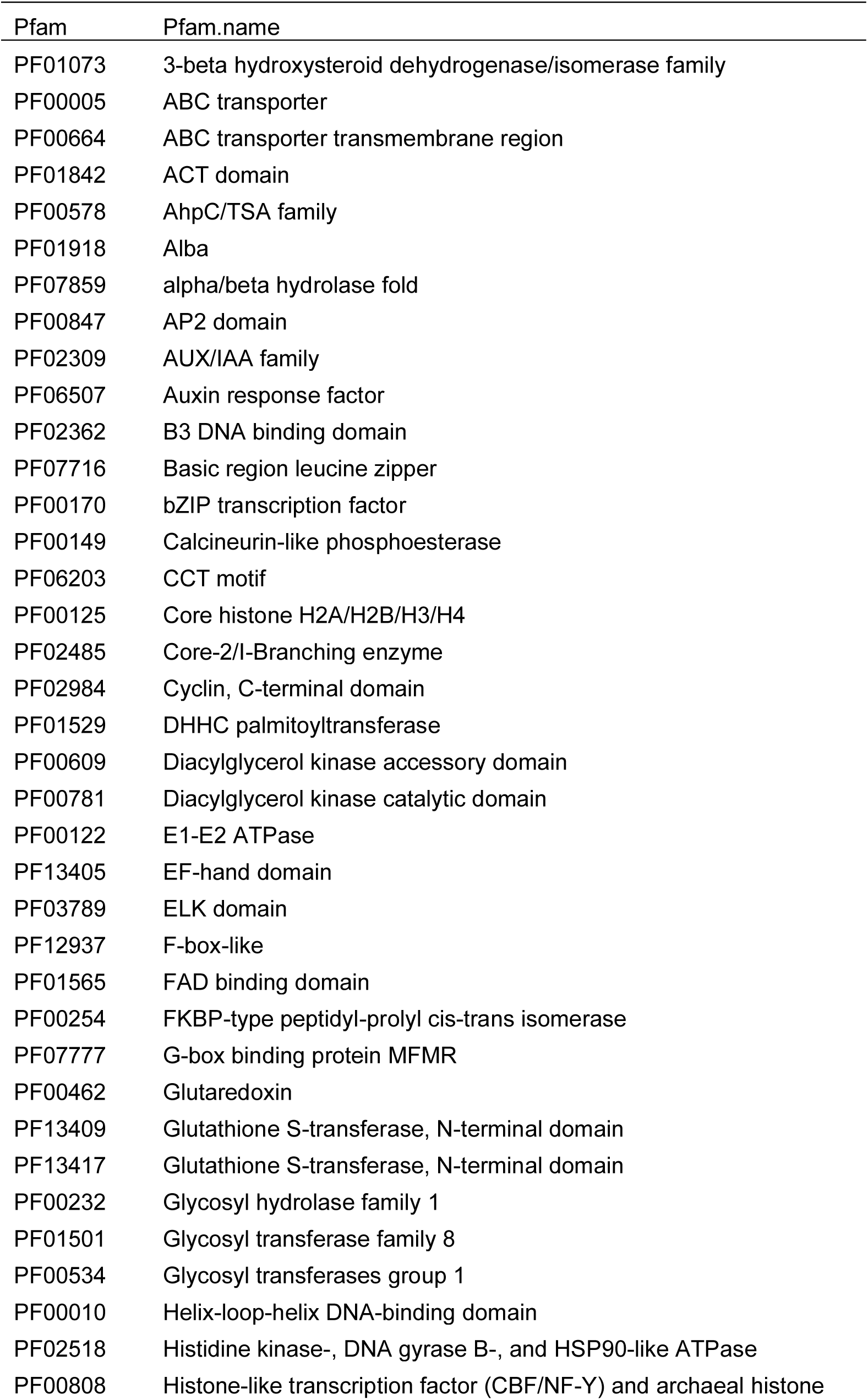
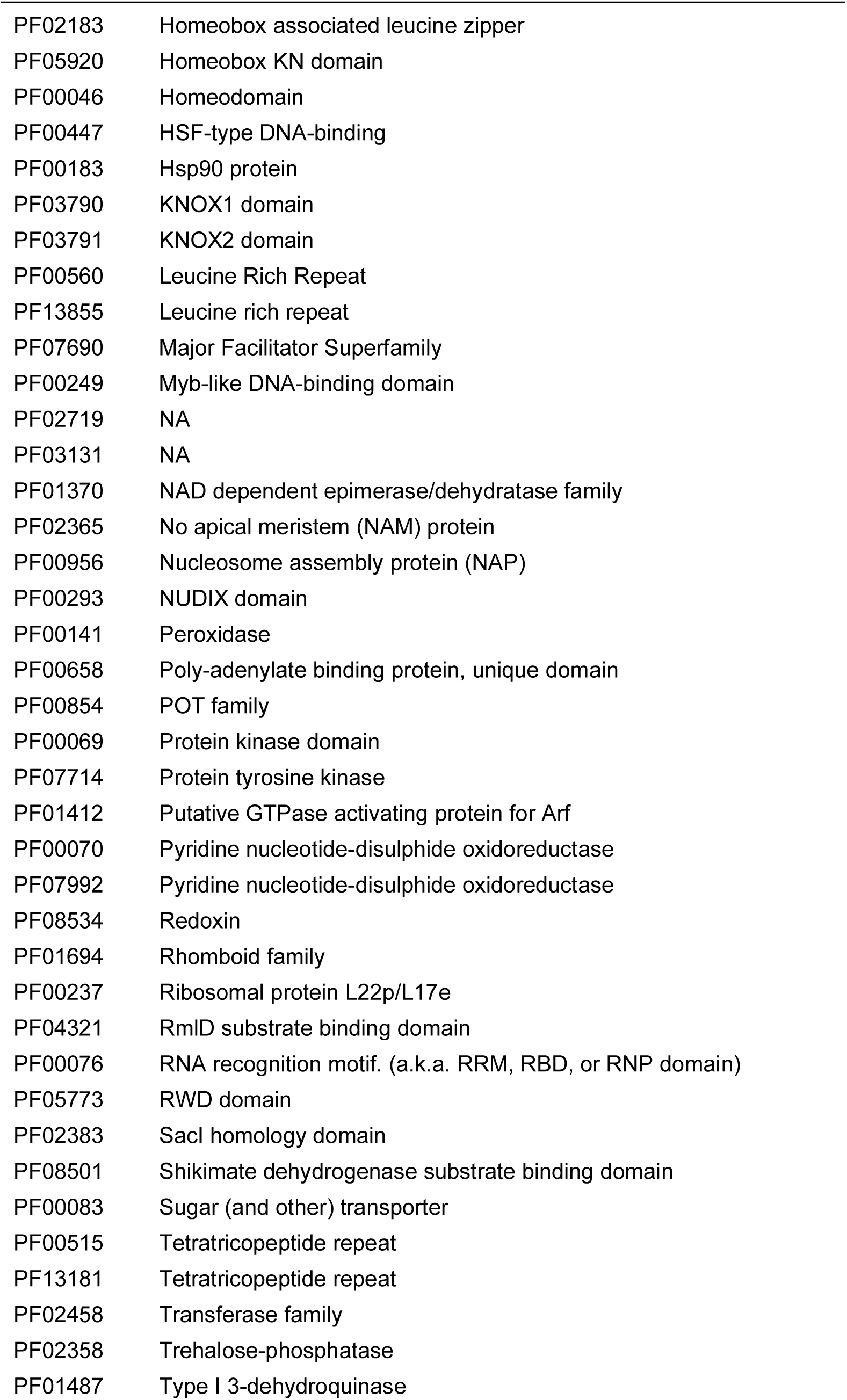
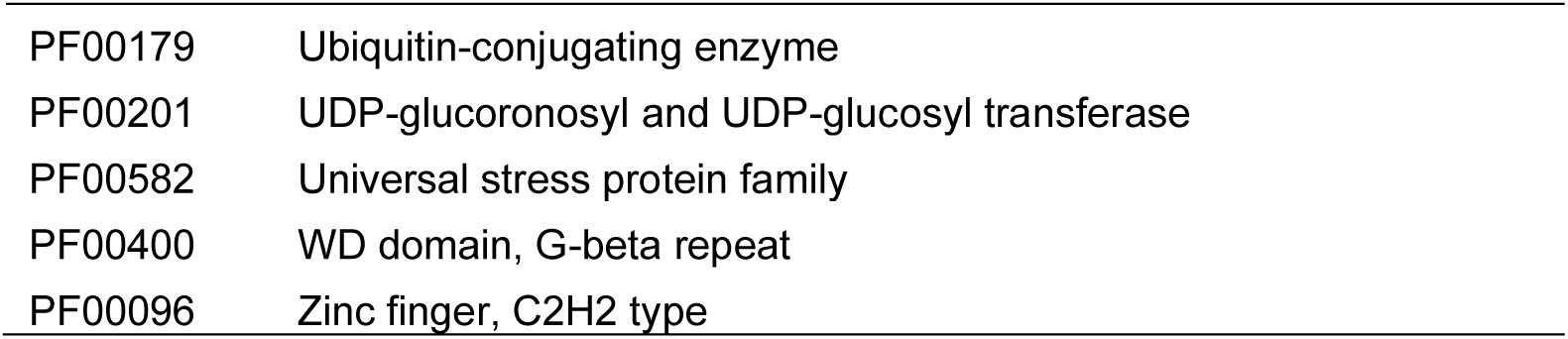
Protein families (Pfams), which were differentially expressed in two drought studies (Gugger et al., 2017, Mead et al., 2019).

| Pfam | Pfam.name |
| --- | --- |
| PF01073 | 3-beta hydroxysteroid dehydrogenase/isomerase family |
| PF00005 | ABC transporter |
| PF00664 | ABC transporter transmembrane region |
| PF01842 | ACT domain |
| PF00578 | AhpC/TSA family |
| PF01918 | Alba |
| PF07859 | alpha/beta hydrolase fold |
| PF00847 | AP2 domain |
| PF02309 | AUX/IAA family |
| PF06507 | Auxin response factor |
| PF02362 | B3 DNA binding domain |
| PF07716 | Basic region leucine zipper |
| PF00170 | bZIP transcription factor |
| PF00149 | Calcineurin-like phosphoesterase |
| PF06203 | CCT motif |
| PF00125 | Core histone H2A/H2B/H3/H4 |
| PF02485 | Core-2/l-Branching enzyme |
| PF02984 | Cyclin, C-terminal domain |
| PF01529 | DHHC palmitoyltransferase |
| PF00609 | Diacylglycerol kinase accessory domain |
| PF00781 | Diacylglycerol kinase catalytic domain |
| PF00122 | E1-E2 ATPase |
| PF13405 | EF-hand domain |
| PF03789 | ELK domain |
| PF12937 | F-box-like |
| PF01565 | FAD binding domain |
| PF00254 | FKBP-type peptidyl-prolyl cis-trans isomerase |
| PF07777 | G-box binding protein MFMR |
| PF00462 | Glutaredoxin |
| PF13409 | Glutathione S-transferase, N-terminal domain |
| PF13417 | Glutathione S-transferase, N-terminal domain |
| PF00232 | Glycosyl hydrolase family 1 |
| PF01501 | Glycosyl transferase family 8 |
| PF00534 | Glycosyl transferases group 1 |
| PF00010 | Helix-loop-helix DNA-binding domain |
| PF02518 | Histidine kinase-, DNA gyrase B-, and HSP90-like ATPase |
| PF00808 | Histone-like transcription factor (CBF/NF-Y) and archaeal histone |
| PF02183 | Homeobox associated leucine zipper |
| PF05920 | Homeobox KN domain |
| PF00046 | Homeodomain |
| PF00447 | HSF-type DNA-binding |
| PF00183 | Hsp90 protein |
| PF03790 | KNOX1 domain |
| PF03791 | KNOX2 domain |
| PF00560 | Leucine Rich Repeat |
| PF13855 | Leucine rich repeat |
| PF07690 | Major Facilitator Superfamily |
| PF00249 | Myb-like DNA-binding domain |
| PF02719 | NA |
| PF03131 | NA |
| PF01370 | NAD dependent epimerase/dehydratase family |
| PF02365 | No apical meristem (NAM) protein |
| PF00956 | Nucleosome assembly protein (NAP) |
| PF00293 | NUDIX domain |
| PF00141 | Peroxidase |
| PF00658 | Poly-adenylate binding protein, unique domain |
| PF00854 | POT family |
| PF00069 | Protein kinase domain |
| PF07714 | Protein tyrosine kinase |
| PF01412 | Putative GTPase activating protein for Arf |
| PF00070 | Pyridine nucleotide-disulphide oxidoreductase |
| PF07992 | Pyridine nucleotide-disulphide oxidoreductase |
| PF08534 | Redoxin |
| PF01694 | Rhomboid family |
| PF00237 | Ribosomal protein L22p/L17e |
| PF04321 | RmID substrate binding domain |
| PF00076 | RNA recognition motif. (a.k.a. RRM, RBD, or RNP domain) |
| PF05773 | RWD domain |
| PF02383 | SacI homology domain |
| PF08501 | Shikimate dehydrogenase substrate binding domain |
| PF00083 | Sugar (and other) transporter |
| PF00515 | Tetratricopeptide repeat |
| PF13181 | Tetratricopeptide repeat |
| PF02458 | Transferase family |
| PF02358 | Trehalose-phosphatase |
| PF01487 | Type I 3-dehydroquinase |
| PF00179 | Ubiquitin-conjugating enzyme |
| PF00201 | UDP-glucuronosyl and UDP-glucosyl transferase |
| PF00582 | Universal stress protein family |
| PF00400 | WD domain, G-beta repeat |
| PF00096 | Zinc finger, C2H2 type |

